# Structural basis of target DNA recognition by CRISPR-Cas12k for RNA-guided DNA transposition

**DOI:** 10.1101/2021.07.07.451486

**Authors:** Renjian Xiao, Shukun Wang, Ruijie Han, Zhuang Li, Clinton Gabel, Indranil Arun Mukherjee, Leifu Chang

## Abstract

The type V-K CRISPR-Cas system, featured by Cas12k effector with a naturally inactivated RuvC domain and associated with Tn7-like transposon for RNA-guided DNA transposition, is a promising tool for precise DNA insertion. To reveal the mechanism underlying target DNA recognition, we determined a cryo-EM structure of Cas12k from cyanobacteria *Scytonema hofmanni* in complex with a single guide RNA (sgRNA) and a double-stranded target DNA. Coupled with mutagenesis and *in vitro* DNA transposition assay, our results revealed mechanisms for the recognition of the GGTT PAM sequence and the structural elements of Cas12k critical for RNA-guided DNA transposition. These structural and mechanistic insights should aid in the development of type V-K CRISPR-transposon systems as tools for genome editing.

## INTRODUCTION

The Clustered Regularly Interspaced Short Palindromic Repeats (CRISPR) and CRISPR-associated (Cas) systems are adaptive immunity systems in bacteria and archaea against mobile genetic elements (MGEs) and have been developed as tools for genome editing (Mohanraju et al., 2016; Sorek et al., 2013). These systems employ guide RNAs and effector proteins to specifically target MGEs for degradation. The arms race between prokaryotes and foreign MGEs has resulted in diverse CRISPR-Cas systems, which are divided into two classes (1 and 2) and six different types (I–VI) (Makarova et al., 2015; Shmakov et al., 2015). The class 2 type V system, featured by a conserved C-terminal RuvC nuclease domain in their effector Cas12 proteins, is abundant and further classified into 11 subtypes (V-A to V-K) (Makarova et al., 2020; Yan et al., 2019), which are promising for the development of new genome editing tools. Among them, Cas12a, Cas12b, and Cas12e have been successfully applied in genome editing (Liu et al., 2019; Strecker et al., 2019a; Teng et al., 2018; Zetsche et al., 2015), such as gene knock-out utilizing non-homologous end joining (NHEJ) to gene knock-in using homology-directed repair (HDR) (Moreno-Mateos et al., 2017). However, the CRISPR-based gene knock-in in mammalian cells relies heavily on endogenous HDR during the S/G2 phase of the cell cycle (Gratz et al., 2014; Moreno-Mateos et al., 2017). Recent discoveries in transposon-associated CRISPR-Cas systems have shed light on the development of an efficient knock-in method independent of host DNA repair pathways, including the type V-K, type I-F, and type I-B systems (Faure et al., 2019; Klompe et al., 2019; Saito et al., 2021; Strecker et al., 2019b). Among those systems, the type V-K system has the advantage of a small effector protein that is more amenable for delivery into mammalian cells.

Cas12k, the effector protein of the type V-K CRISPR-Cas system, was first identified with a featured inactive RuvC nuclease domain, which led to the subsequent discovery of its association with Tn7-like transposons (Faure et al., 2019; Strecker et al., 2019b). Strecker *et al* showed that a CRISPR-associated transposase from cyanobacteria *Scytonema hofmanni* (ShCAST) can be directed and inserted to target sites 60 to 66 base pairs downstream of the protospacer adjacent motif (PAM). The ShCAST system contains a CRISPR module composed of Cas12k with a CRISPR array and a transposon module **(Fig. S1A)**. The transposon module contains a single component transposase TnsB (similar to MuA transposase in the transposable phage Mu (Montano et al., 2012)), an AAA+ regulator TnsC, a target selector TniQ (homologue of TnsD in Tn7 transposon (Faure et al., 2019)), and the left and right ends of the transposon. This system was successfully repurposed for efficient RNA-guided DNA insertion in *E. coli* (Strecker et al., 2019b). However, further development of this tool for genome editing, biomedical applications, and the eventual treatment of human diseases requires a deeper understanding of the molecular mechanism underlying RNA-guided DNA transposition.

Here we report the cryo-EM structures of a Cas12k–sgRNA–target DNA ternary complex and a Cas12k–sgRNA binary complex. The structures, combined with *in vitro* transposition assay, provide mechanistic insights into target recognition and RNA-guided DNA transposition by the ShCAST system.

## RESULTS

### *In vitro* DNA transposition

We first purified Cas12k, sgRNA and transposition proteins (TnsB, TnsC and TniQ) in the ShCAST system and tested their function using a previously established *in vitro* DNA transposition assay (Strecker et al., 2019b) **(Fig. S1B,C)**. Our results suggest that TnsB, TnsC, and magnesium (required for transposon end cleavage and target joining (Skelding et al., 2002)) are strictly required for DNA transposition; whereas additional components including Cas12k, sgRNA, and TniQ are necessary for RNA-guided DNA transposition **(Fig. S1D–G)**. Omitting any of the later three components leads to DNA transposition in a non-RNA-guided manner. Out of ten randomly selected colonies from the assay using all components, eight RNA-guided and two non-RNA-guided insertions were observed **(Fig. S1G)**. Both simple insertion and co-integration products were observed in RNA-guided insertions (Rice et al., 2020; Strecker et al., 2020), with co-integration being the major product in our experiments as revealed by restriction enzyme digestion and DNA sequencing **(Fig. S1C,H,I)**.

### Overall structure of Cas12k–sgRNA–target DNA

To understand the mechanism of RNA-guided target DNA recognition, we assembled a Cas12k–sgRNA–target DNA ternary complex by incubating Cas12k, sgRNA, and a target DNA containing a GGTT PAM sequence **(Fig. S1J, and Table S1)**. Using cryo-EM, we reconstructed a map of this ternary complex at 3.6 Å resolution **(Figs. 1B,C and S2, and Table 1)**, which allowed us to build the atomic model **(Fig. S3A)** except residues 103–270 of Cas12k, the crRNA–target DNA heteroduplex beyond 10 bp from the PAM duplex, and small regions (e.g. 1–8 nt of sgRNA), which are not resolved in the map most likely due to flexibility.

**Figure 1.**
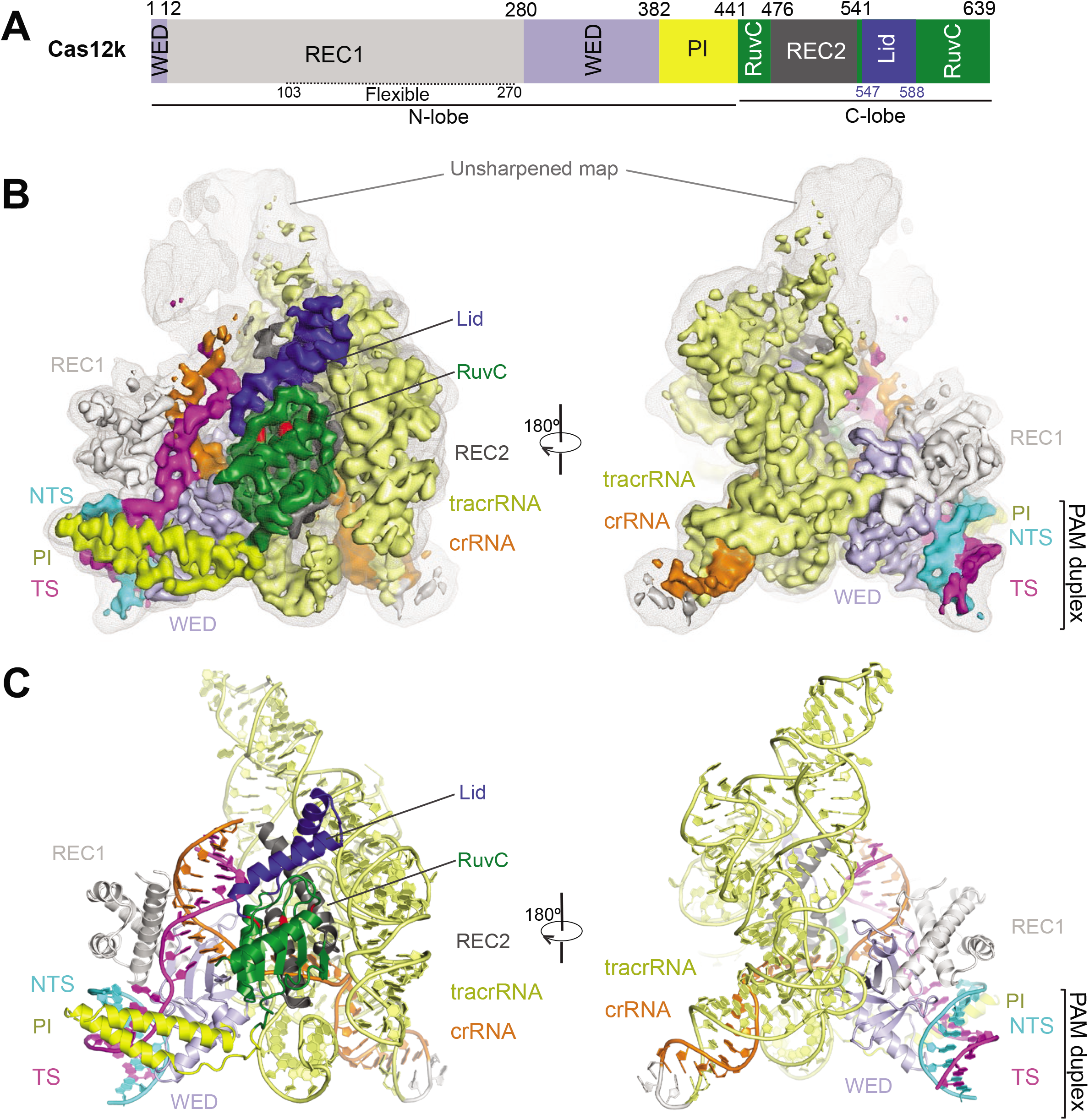
Overall structure of the Cas12k–sgRNA–target DNA complex. **(A)** Schematic of domain organization of Cas12k based on structure. **(B)** Cryo-EM map of the Cas12k–sgRNA– target DNA complex at 3.6 Å in two views with each domain color coded as in **A**. The unsharpened map is shown in grey mesh. **(C)** Atomic model of the Cas12k–sgRNA–target DNA complex shown in cartoon in the same views as in **B**.

**Table 1.**
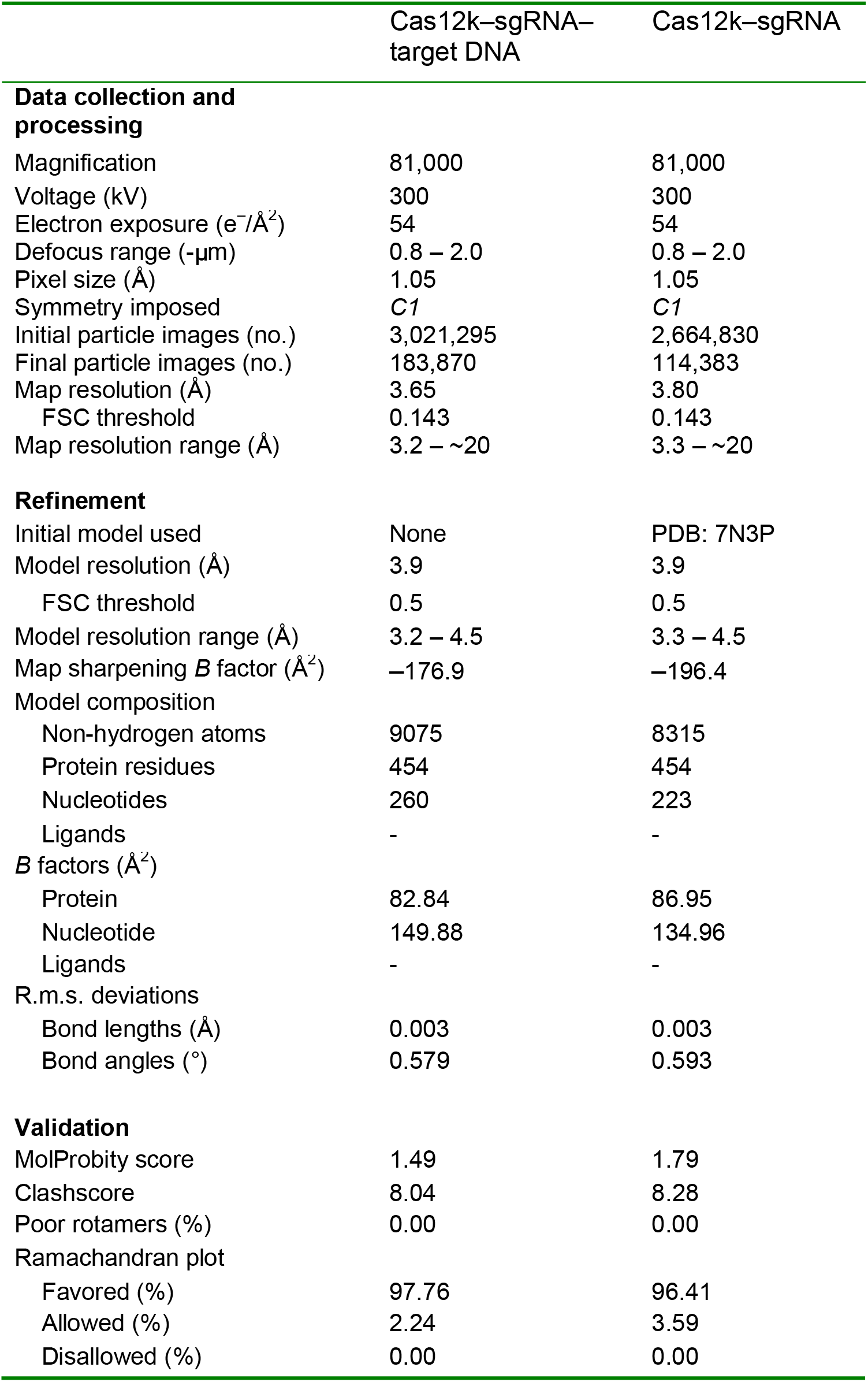
Cryo-EM data collection, refinement and validation statistics.

The overall structure of Cas12k resembles other Cas12 proteins, with Cas12f as the closest match by a DALI search (z-score, 14.5) (Takeda et al., 2020; Xiao et al., 2021) **(Fig. 2A,B)**. The 637-residue protein adopts a bi-lobed structure connected by a loop. The N-terminal lobe of Cas12k is composed of the WED, REC1, and PI domains. The WED domain, which plays a major role in recognizing sgRNA, contains seven strands (β1–7) with a helix α5 inserted between β5 and β6 **(Fig. 2C)**. The REC1 domain is inserted between β1 and β2 of the WED domain and composed of an N-terminal helical bundle α1–4 (REC1^13–102^) and a C-terminal flexible region (REC1^103–270^) that is predicted to form 6–7 helices **(Figs. 2D and S3B)**. Although sharing low sequence similarity, REC1^13–102^ is structurally similar to the REC1^C^ domain in Cas12f **(Figs. 2D and S3B,C)**, which forms the dimerization interface in Cas12f (Takeda et al., 2020; Xiao et al., 2021). However, the key hydrophobic residues (I118, Y121, Y122, I126) in Cas12f are not conserved in REC1^13–102^ **(Figs. 2D and S3B)**, which may be a reason why a Cas12k dimer is not observed. Following the WED domain is a PI domain composed of two helices, α6 and α7, which is absent in Cas12f but observed in some other Cas12 proteins such as Cas12i (Zhang et al., 2020).

**Figure 2.**
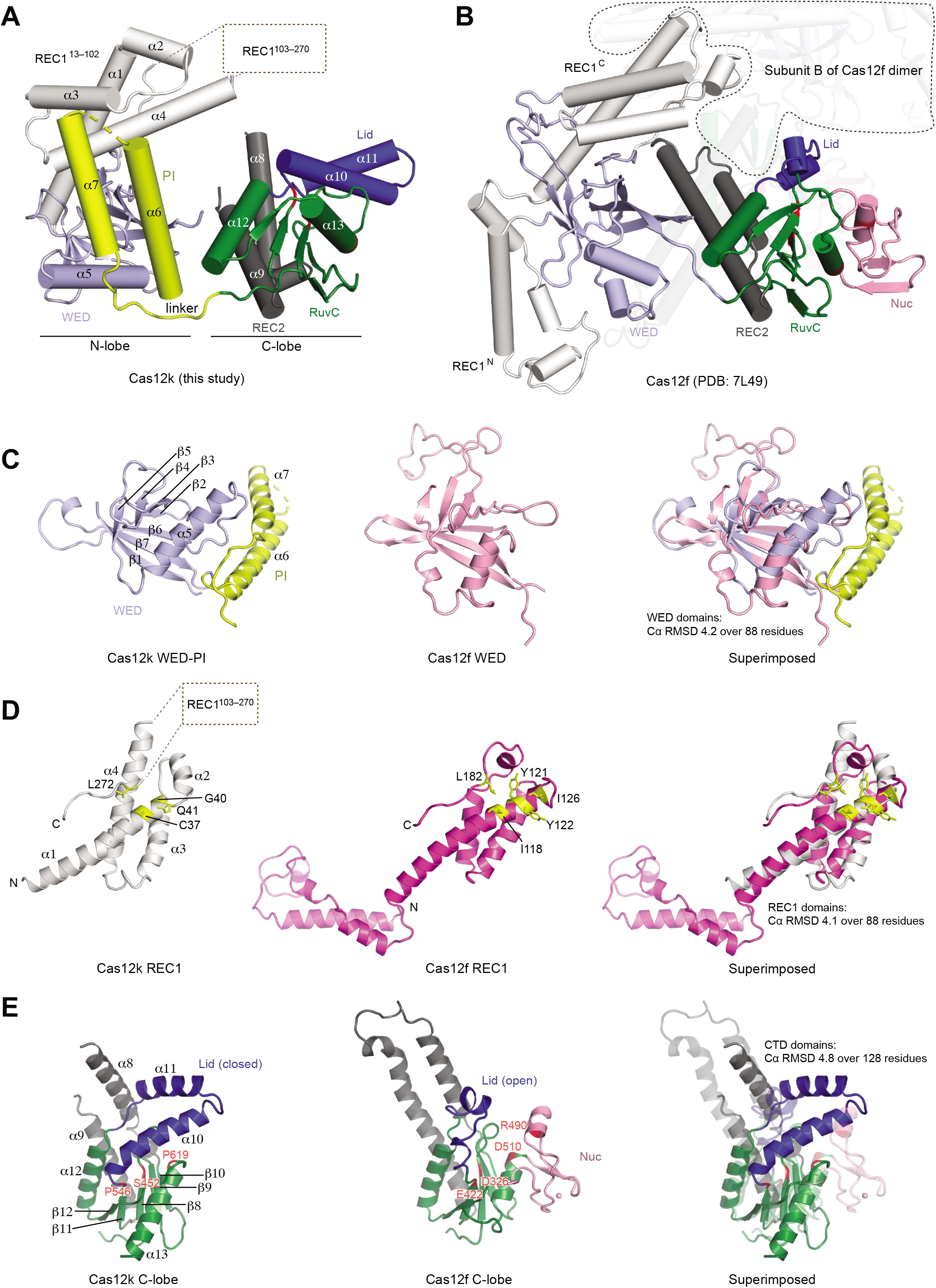
Structure of Cas12k and comparison with Cas12f. **(A,B)** Atomic models of Cas12k (**A**) and Cas12f (PDB: 7L49) (**B**). The subunit A of Cas12f is shown in the same view as Cas12k whereas the subunit B is semi-transparent. **(C–E)** Comparison of the domains between Cas12k and Cas12f.

The C-terminal lobe of Cas12k is composed of the RuvC and REC2 domains. Both the sequence and structure of the RuvC domain of Cas12k are conserved relative to Cas12f; however, the Cas12f’s triplet of acidic residues required for nuclease activity is replaced by either serine or proline in Cas12k **(Fig. S3B,C)**. A Cas12k mutant restoring the catalytic residues (S452D, P546E, and P619D) did not reinstate target DNA cleavage (**Fig. S1K**). In addition to the altered catalytic residues, two additional features are observed in the RuvC domain of Cas12k compared to that of Cas12f. First, there is no Nuc domain in Cas12k **(Figs. 2A,B,E and S3B)**. The Nuc domains or equivalent domains are inevitably present in all Cas12 proteins with structures determined to date including Cas12a (Dong et al., 2016; Gao et al., 2016; Nishimasu et al., 2017; Stella et al., 2017; Stella et al., 2018b; Swarts and Jinek, 2019; Swarts et al., 2017; Yamano et al., 2016; Yamano et al., 2017; Zhang et al., 2019), Cas12b (Liu et al., 2017; Wu et al., 2017; Yang et al., 2016), Cas12e (Liu et al., 2019), Cas12f (Takeda et al., 2020; Xiao et al., 2021), Cas12g (Li et al., 2021) and Cas12i (Huang et al., 2020; Zhang et al., 2020), and play an essential role in the nuclease activity. Second, the lid motif of Cas12k is longer than that of Cas12f and is in a closed conformation that covers the pseudonuclease site **(Fig. 2A,C)**. Both features are consistent with the lack of nuclease activity in the RuvC domain of Cas12k. Taken together, Cas12k is closely related to Cas12f in structure but may have evolved these new features to meet the requirements for DNA transposition.

### Structure of sgRNA

The 265-nt sgRNA is composed of a 44-nt crRNA (30-nt spacer and 14-nt repeat) and a 218-nt tracrRNA connected by a short 3-nt linker **(Fig. 3A,B and Table S1)**. The sgRNA contains two anti-repeat:repeat (AR:R) duplexes, AR:R1 and AR:R2, formed by the repeat region of crRNA (−1 to −5 & −6 to −14) and the anti-repeat sequences of tracrRNA (87–91 & 211–218). The AR:R1 duplex, clamped by the WED and REC2 domains, is the major region on the sgRNA that bridges the N- and C-lobes of Cas12k **(Figs. 3C and S4A)**. The AR:R2 duplex makes no contact with Cas12k, and could function mainly for the recognition between crRNA and tracrRNA **(Fig. 1C)**.

**Figure 3.**
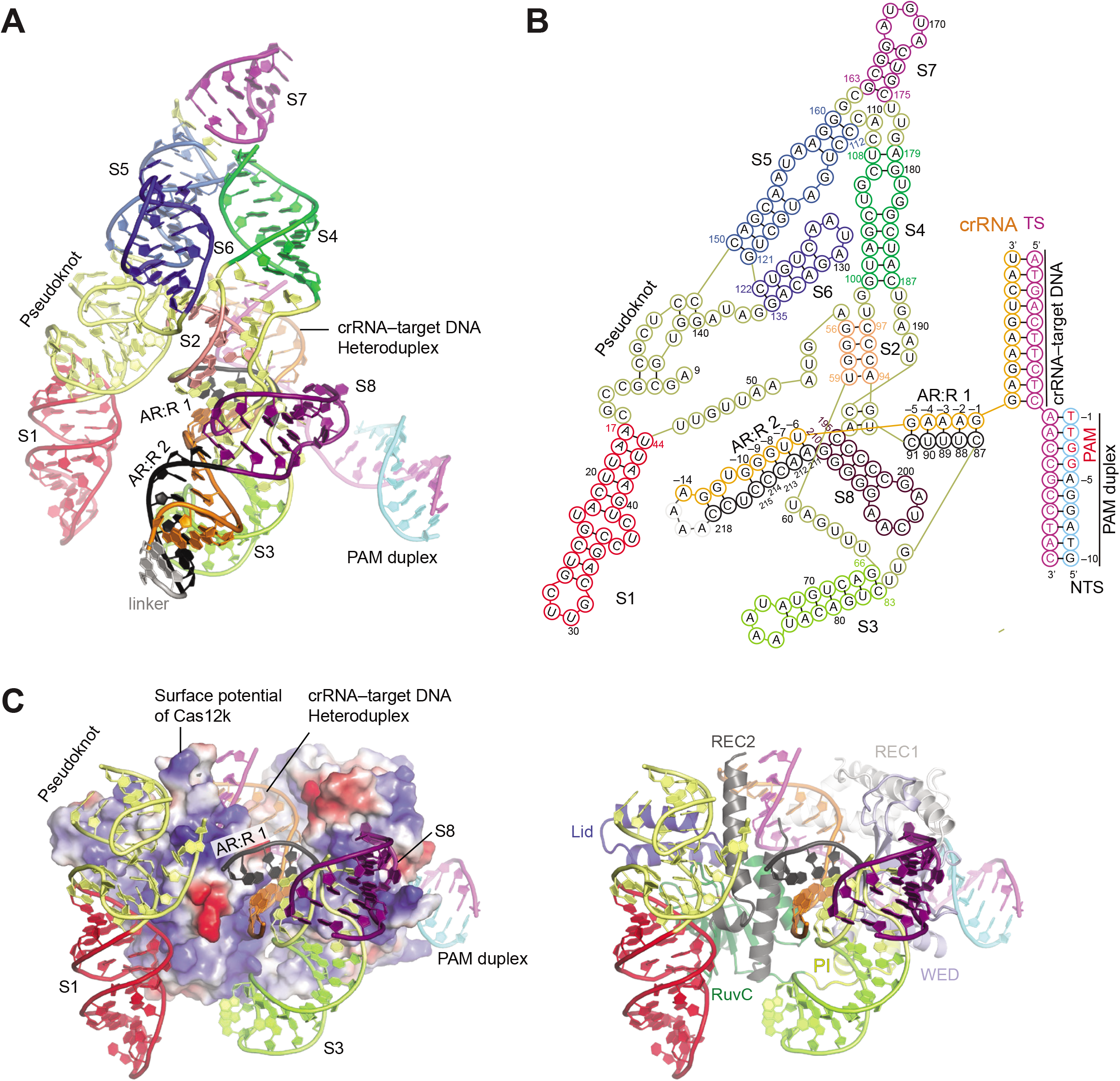
Overall structure of sgRNA. **(A)** Structure of the sgRNA and target DNA in the Cas12k–sgRNA–target DNA complex in cartoon presentation with stem-loops (S1–8), pseudoknot, and AR:R 1–2 duplexes color coded. **(B)** Schematic of the sgRNA and target DNA, color coded as in **A**. **(C)** Contacts between sgRNA and Cas12k. Stem loops not in contact with Cas12k are not shown. Cas12k in surface potential and cartoon representations are shown in left and right panels, respectively.

The tracrRNA part (1–218) contains five stem loops formed by local regions including S1 (17– 44), S3 (66–83), S6 (122–135), S7 (163–175), and S8 (195–210) and three duplexes formed by long-distance regions such as S2 (56–59 & 94–97), S4 (100–108 & 179–187), and S5 (112–121 & 150–160) **(Fig. 3A,B)**. Between S1 and S2 is a pseudoknot structure formed by three fragments (9–16, 45–55, 140–149). S1 and the pseudoknot contact the C-lobe of Cas12k **(Figs. 3C and S4B)**, whereas S3 and S8 interact with the N-lobe of Cas12k primarily through electrostatic interactions **(Figs. 3C and S4C)**. A previous study showed that deletion of 1–47 nt in the sgRNA completely abolished RNA-guided DNA insertion, indicating that S1 is essential (Strecker et al., 2019b). S4–7 show no contact with Cas12k and display considerate flexibility revealed by 3D variance analysis **(Fig. S2D–F)**.

### PAM recognition

The PAM duplex is enclosed in a positively charged groove formed by the REC1, WED, and PI domains **(Fig. 4A).** All three domains contribute residues that directly interact with the bases of the PAM sequence for sequence-specific recognition **(Figs. 4B,C and S4D,E)**. Specifically, R78 from the REC1 domain establishes two hydrogen bonds with the base G(−3) of the non-target strand (NTS). The hydroxyl group of T287 from the WED domain forms two hydrogen bonds with base A(−2) of the target strand (TS). R421 from the PI domain interacts with both A(−2) and T(−1) from the TS and NTS, respectively. In addition, a number of polar or positively charged residues recognize the PAM duplex through the phosphate backbones **(Figs. 4C and S4D,E)**. To test the structural observations, we mutated the three residues that recognize the bases and two positively charged residues that bind to phosphate backbones, R350 and R428 from the WED and PI domains, respectively. Alanine substitution of any of the residues reduced *in vitro* RNA-guided transposition activity by PCR readout (**Figs. 4D, S1E–G and S5**).

**Figure 4.**
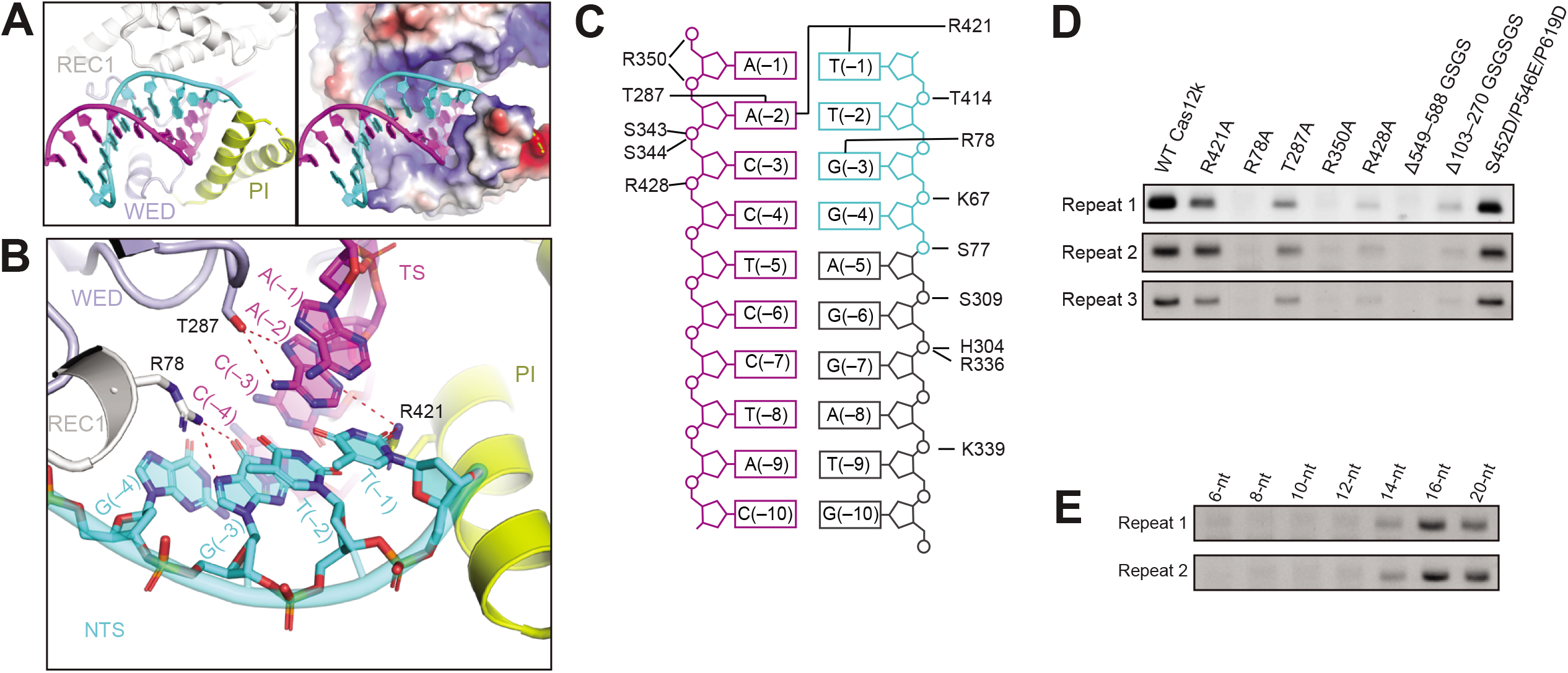
Target DNA recognition by Cas12k. **(A)** The PAM duplex is bound to a positively charged grove by REC1, WED, and PI domains (shown in cartoon and surface potential in the left and right panels, respectively). **(B)** Detailed interactions between the PAM duplex and Cas12k. Interactions are indicated by red dashed lines. **(C)** Schematic of the interactions between the PAM duplex and Cas12k. **(D)** PCR results of *in vitro* DNA transposition assay using wild-type Cas12k and various Cas12k mutants. The results shown are from three replicates. **(E)** PCR results of *in vitro* DNA transposition assay using sgRNA with different spacer length. The results shown are from two replicates.

### crRNA-DNA heteroduplex recognition

A 10-bp crRNA–target DNA heteroduplex is observed in the ternary complex, which primarily contacts the REC1 and REC2 domains, as well as the lid motif **(Figs. 3C and S4F,G)**. The heteroduplex beyond 10 bp is likely formed but in a flexible state, as it is visible in the unsharpened map at a lower contour level **(Fig. 1B).** This shorter stabilized heteroduplex is in contrast to the usual 20-bp heteroduplex observed in other Cas12 structures where the PAM distal end of the heteroduplex is stabilized by either relatively large REC1 and REC2 domains (e.g. Cas12a (Dong et al., 2016; Gao et al., 2016; Nishimasu et al., 2017; Stella et al., 2017; Stella et al., 2018b; Swarts and Jinek, 2019; Swarts et al., 2017; Yamano et al., 2016; Yamano et al., 2017; Zhang et al., 2019), Cas12b (Liu et al., 2017; Wu et al., 2017; Yang et al., 2016), Cas12e (Liu et al., 2019) and Cas12i (Huang et al., 2020; Zhang et al., 2020)) or a second protein subunit (e.g. Cas12f (Takeda et al., 2020; Xiao et al., 2021)). The flexible REC1^103–270^ might provide additional interactions with the heteroduplex. Cas12k with REC1^103–270^ deleted (Δ103–270GSGSGS) is inactive for RNA-guided DNA transposition **(Figs. 4D and S5)**.

Guided by our observation of a shorter stabilized heteroduplex, we set out to determine the minimum spacer length in sgRNA required for RNA-guided DNA transposition. We designed sgRNA with various spacer length, including 6, 8, 10, 12, 14, 16, 18, and 20 nucleotides **(Table S1)**. Our *in vitro* DNA transposition assay suggests that at least a 14-nt spacer length is required for detectable DNA transposition, and at least 16-nt is required for optimized activity **(Fig. 4E)**. This is consistent with a recent study showing that 16 nt is both sufficient and near the minimum length required for insertion in the type V-K system (Saito et al., 2021). This result suggests that one checkpoint for transposition in the type V-K system is the formation of the crRNA-target DNA heteroduplex at 14–16 bp.

### Conformational changes induced by target DNA recognition

To understand the conformational changes in Cas12k upon target DNA recognition, we reconstructed a cryo-EM structure of the Cas12k–sgRNA binary complex at 3.8 Å **(Figs. 5A,B and S6, and Table 1)**. Although still largely flexible, REC1^103–270^ is more visible in the binary complex and contacts the lid motif in the RuvC domain **(Fig. 5C)**. Structural superimposition with the ternary structure reveal minimal conformational change in Cas12k, with the exception of the REC1 domain that undergoes a ~6–8 Å shift to contact target DNA **(Fig. 5D,E)**. To be noted, the lid motif adopts a similar closed conformation in both the ternary and binary complex. The closed-to-open transition of the lid motif upon target DNA recognition is shown as a conserved mechanism for activation of the RuvC nuclease activity in Cas12 proteins (Stella et al., 2018a; Xiao et al., 2021; Zhang et al., 2020). However, deletion of the lid motif (Δ549–588GSGS) completely abolishes RNA-guided DNA transposition **(Figs. 4D and S5)**, suggesting it plays an essential function.

**Figure 5.**
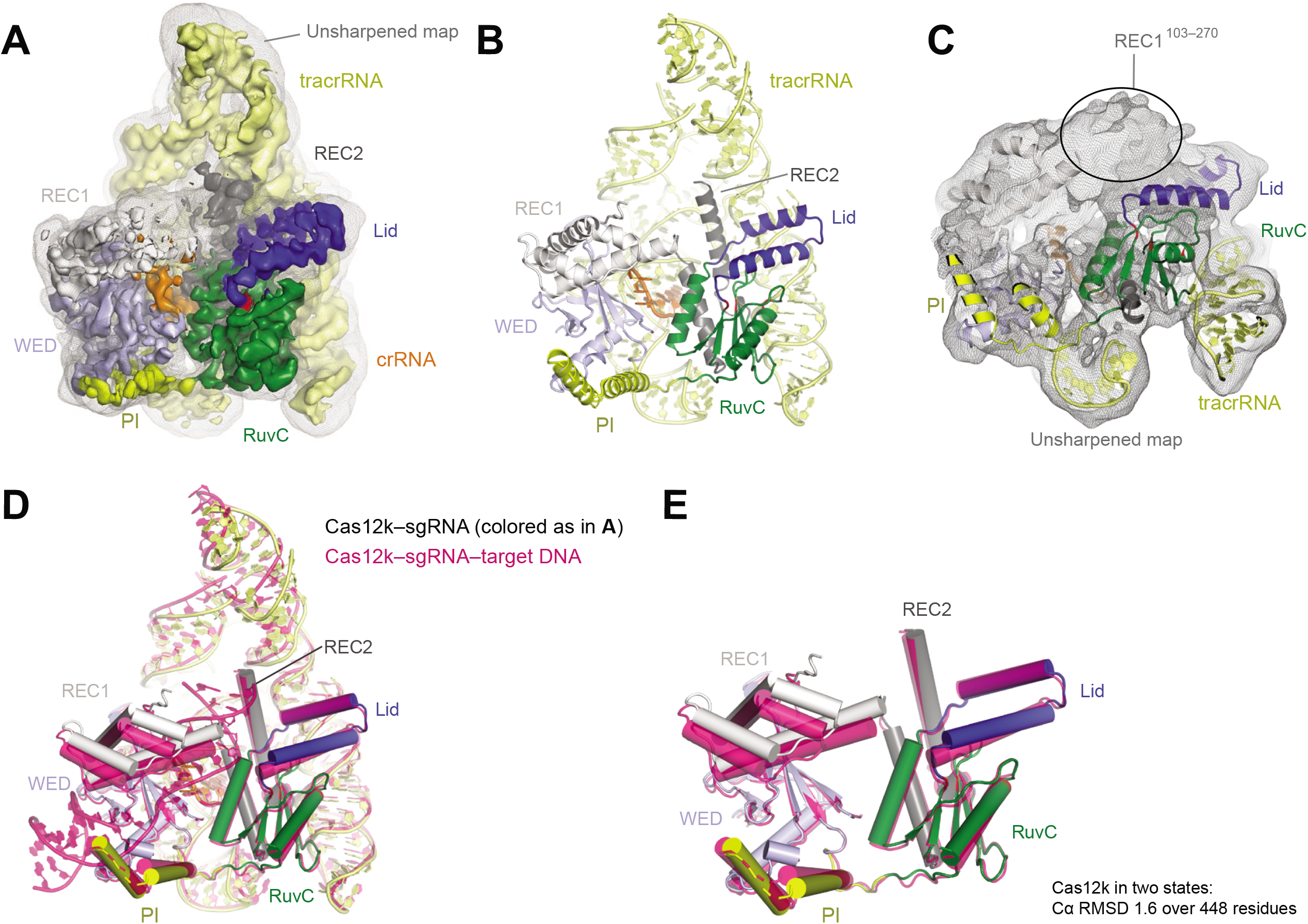
Structure of the Cas12k-sgRNA complex. **(A)** Cryo-EM map of the Cas12k–sgRNA complex at 3.8 Å with each subunit color coded as in **Fig. 1A**. The unsharpened map is shown in grey mesh. **(B)** Atomic model of the Cas12k–sgRNA complex shown in cartoon in the same views as in **A. (C)** Cryo-EM density of REC1^103–270^ (circled) in the binary complex. **(D)** Structural superimposition of Cas12k–sgRNA (color-coded as in **A**) and Cas12k–sgRNA–target DNA (magenta) complexes. **(E)** Structural superimposition of Cas12k protein in the two states shown in **D**.

## DISCUSSION

RNA-guided DNA transposition by the type V-K CAST system requires both the CRISPR module and the transposition module (Strecker et al., 2019b). The CRISPR module recognizes target DNA using guide RNAs and recruits the transposition machinery for DNA insertion. In this study, we showed the cryo-EM structure of Cas12k and the mechanism of target DNA recognition by the CRISPR module of the ShCAST system.

Despite sharing similar architecture with other Cas12 proteins, Cas12k displays considerable differences that may be related to its association with the Tn7-like transposon, including an inactive RuvC domain, the absence of the Nuc domain, and a longer and closed lid motif. Although undergoing no closed-to-open conformational change upon recognition of target DNA, the lid motif is kept in Cas12k and essential for RNA-guided DNA transposition. Given the essential function of the lid motif and the lack of nuclease activity of Cas12k, we speculate that the lid motif might play a role in either stabilization of the structure or the recruitment of transposition proteins.

Four stem loops (S4–7) within the sgRNA show no interactions with Cas12k, raising a question about their function. Interestingly, when sgRNA is removed from the *in vitro* transposition assay, the number of colonies are significantly larger compared to that of other conditions **(Fig. S1E)**; however, none of the tested colonies shows RNA-guided DNA insertion. This result may suggest that sgRNA might play an inhibitory role in the transposition machinery for non-RNA-guided DNA insertion. This may not be surprising because to direct the transposon machinery for RNA-guided DNA transposition, the CRISPR-Cas system may have evolved a mechanism to inhibit the transposon’s original activity.

Recent studies showed the role of the AAA+ protein TnsC in transposition target site selection (Park et al., 2021; Shen et al., 2021). In the ShCAST system, TnsC forms filament structure on DNA, which is capped by TniQ (Park et al., 2021). TniQ is likely directly associated to Cas12k, similar to previous observations showing that TniQ is bound to the CRISPR effector complex, the Cascade complex, in the type I-F CAST system (Halpin-Healy et al., 2020; Jia et al., 2020; Li et al., 2020; Wang et al., 2020). The interactions between TnsB transposase and TnsC could direct DNA insertion in a fixed position relative to the target DNA recognition site of the CRIPSR module, which is 60–66 bp downstream of the PAM in the ShCAST system. The Cas12k structure reported here and these recent studies are beginning to unravel the underlying mechanism for RNA-guided DNA transposition.

Transposon-associated CRIPSR-Cas systems are promising tools for gene insertion application; however, possible off-target insertion raises concerns because it can cause genome instability. In the case of the ShCAST system, non-RNA-guided DNA insertion is observed in *in vitro* transposition assay **(Fig. S1G)** and in *E.coli* (Strecker et al., 2019b). To reduce or eliminate this unwanted DNA insertion, further studies will be required to understand detailed mechanisms in the ShCAST system, including the interactions between the Cas12k–sgRNA–target DNA module and the whole transposition machinery; this will be especially vital for manipulating the system in genome editing applications.

## ACKNOWLEDGMENTS

We thank Thomas Klose for help with cryo-EM and Steven Wilson for computation. This work made use of the Purdue Cryo-EM Facility. C.G. is supported by a grant from the NIH [T32GM132024]. This work was supported by the NIH grant R01GM138675 to L.C.

## AUTHOR CONTRIBUTIONS

L.C. supervised the study. R.X., S.W. and R.H. prepared samples. Z.L., R.X., and L.C. collected and processed cryo-EM data. S.W, R.X., and R.H. performed biochemical analysis with help from C.G. and I.A.M. All authors analyzed the data. R.X., S.W., R.H., and L.C prepared the manuscript with input from other authors.

## DATA AVAILABILITY

Cryo-EM reconstructions of Cas12k–sgRNA–target DNA and Cas12k–sgRNA complexes have been deposited in the Electron Microscopy Data Bank under the accession numbers EMD-24143 and EMD-24142, respectively. Coordinates for atomic models of Cas12k–sgRNA–target DNA and Cas12k–sgRNA complexes have been deposited in the Protein Data Bank under the accession numbers 7N3P and 7N3O, respectively.

---

**Figure S1.**
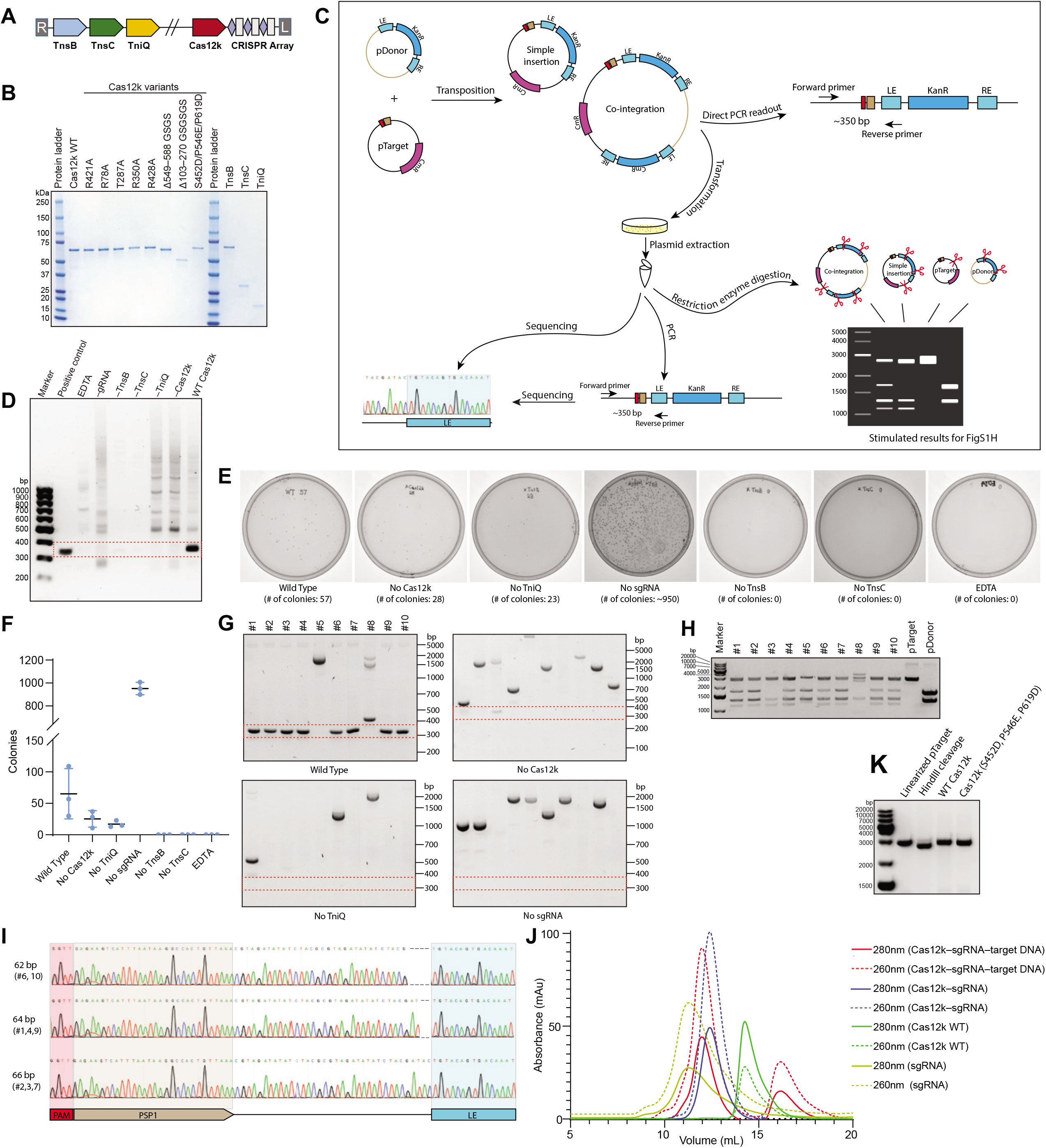
*In vitro* DNA transposition assay. **(A)** Schematic of the genomic organization of the shCAST system. **(B)** SDS-PAGE of purified proteins used for the *in vitro* DNA transposition assay. **(C)** Schematic of the procedures for the *in vitro* DNA transposition assay. **(D)** PCR results using reaction mixture of *in vitro* DNA transposition assay as template. Positions of expected PCR readout at ~350 bp are indicated by a red dashed box. **(E,F)** LB agar plate showing colonies after transformation of each transposition reaction. Graph in **F** shows mean±SD (n=3). **(G)** PCR results using purified plasmid as template. Ten colonies are randomly selected for plasmid extraction from each plate in **E**. Positions of expected PCR readout at ~350 bp are indicated by red dashed boxes. **(H)** Restriction enzyme digestion assay of ten plasmids extracted from the wild type condition in **E**. **(I)** Sanger sequencing of PCR products in the wild type condition in **G**. **(J)** Superimposed size exclusion chromatography profiles of Cas12k samples, including the Cas12k–sgRNA–target DNA ternary complex (red), the Cas12k–sgRNA binary complex (blue), Cas12k (green) and sgRNA (yellow), with UV absorbance curves at 280 nm and 260 nm shown in solid and dashed lines, respectively. **(K)** Target DNA (linearized pTarget plasmid) cleavage assay using wild type Cas12k and Cas12k with the catalytic acidic residues in RuvC restored by mutations. The results shown in **D–E** are representative of more than three experiments.

**Figure S2.**
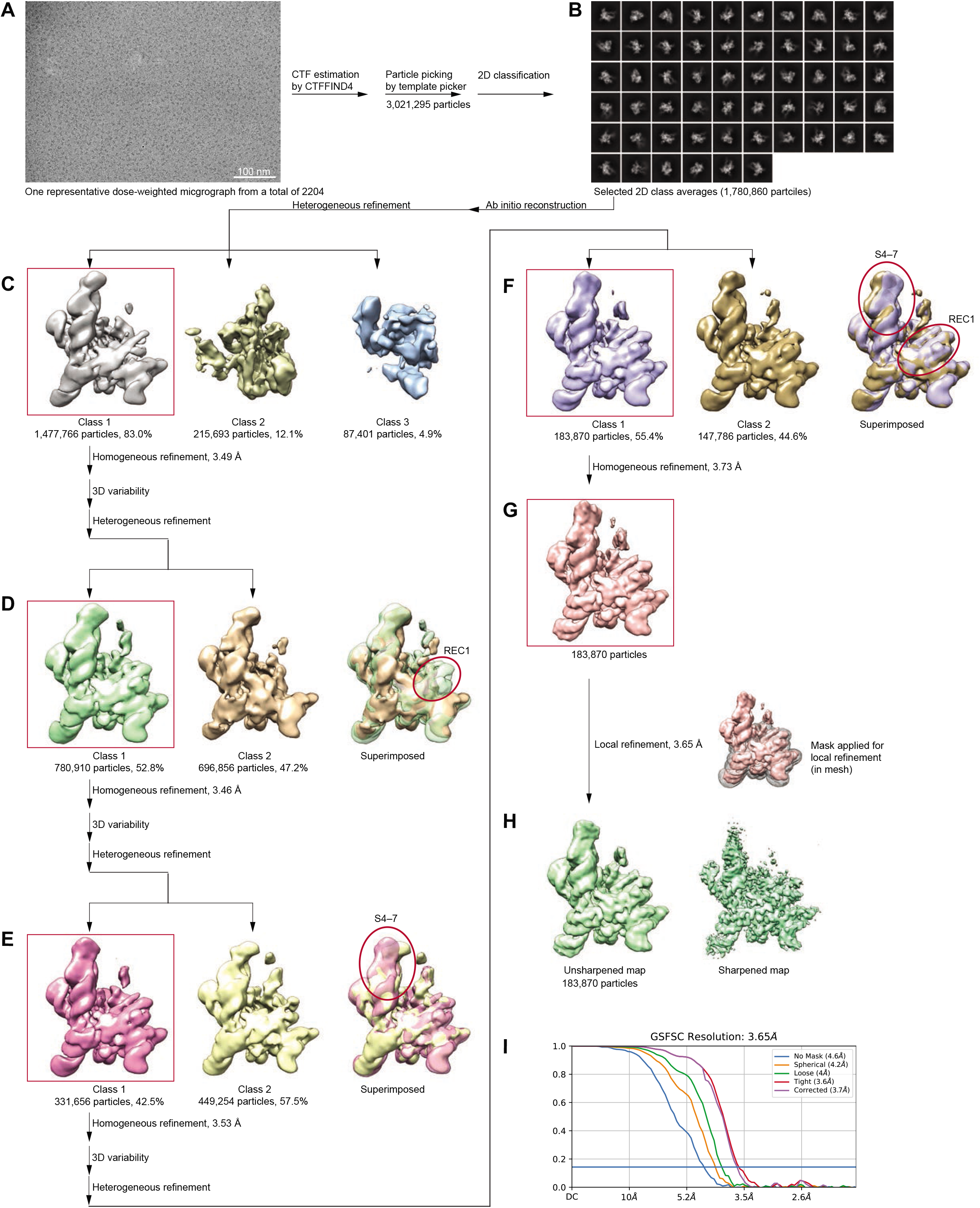
Cryo-EM data processing for the Cas12k–sgRNA–target DNA ternary complex. **(A)** A representative raw cryo-EM micrograph of the Cas12k–sgRNA–target DNA complex from a total of 2204 micrographs. **(B)** Representative, good 2D class averages from a total of 100 images. **(C)** Three 3D reconstructions from heterogeneous refinement. **(D–F)** Three rounds of supervised heterogenous refinement using two maps from 3D variability analysis as templates. Variable regions are indicated by red circles. **(G)** Homogeneous refinement of final particle set. **(H)** Local refinement of final particle set using a mask as indicated. Shown on the left is the unsharpened map, and on the right is the sharpened map. **(I)** Plots of the half-map FSC.

**Figure S3.**
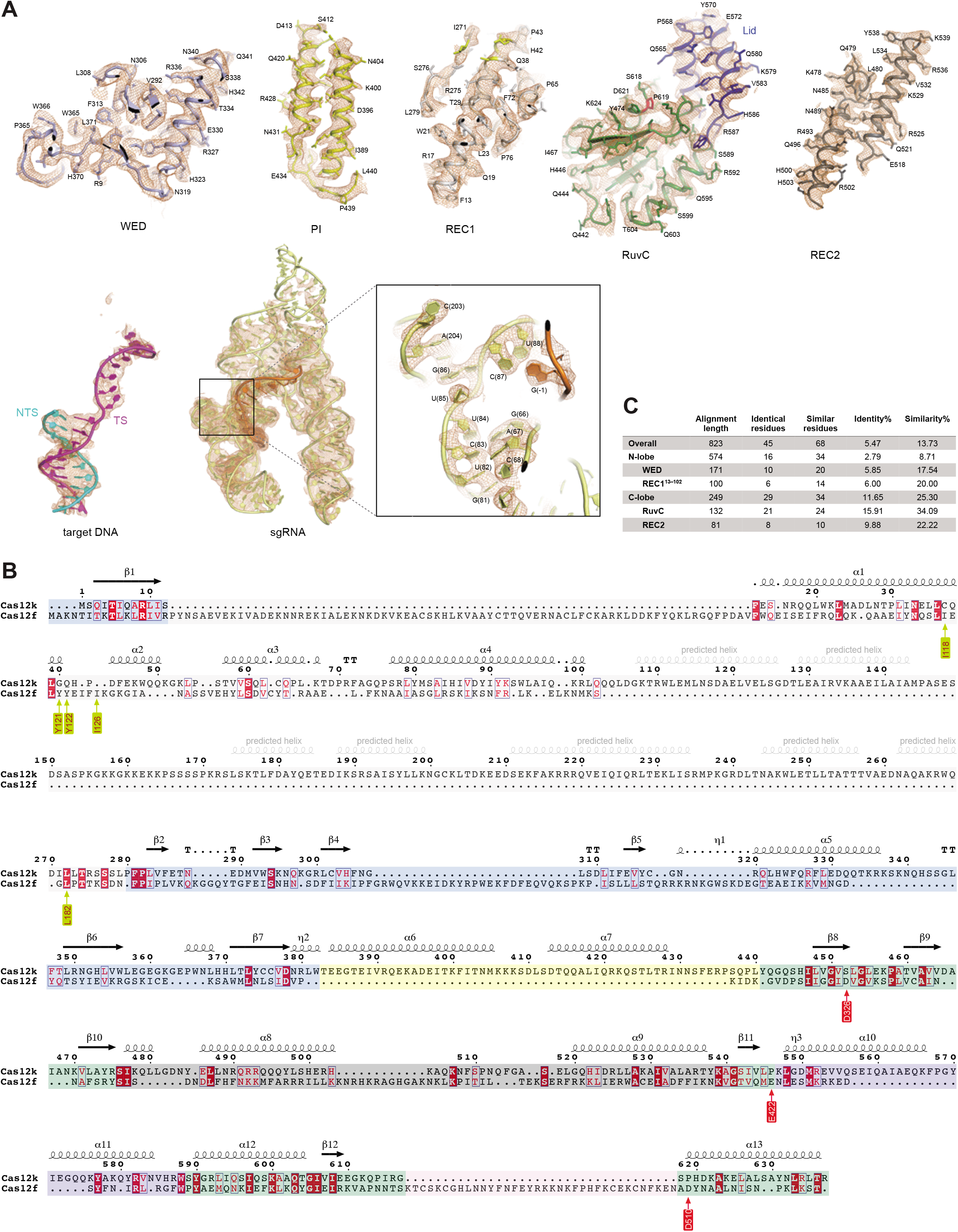
Detailed cryo-EM density map and structure-based sequence alignment. **(A)** Fitting between the cryo-EM map of the Cas12k–sgRNA–target DNA complex and the atomic model. **(B)** Structure based sequence alignment of Cas12k and Cas12f. Residue numbers and secondary structures are labeled according to Cas12k. Arrowed residues are key residues for Cas12f dimerization (in yellow) and catalytic residues in the RuvC domain of Cas12f (in red). Each domain is indicated by background colors as in **Fig. 1A**. **(C)** Sequence identity and similarity based on alignment in **B**.

**Figure S4.**
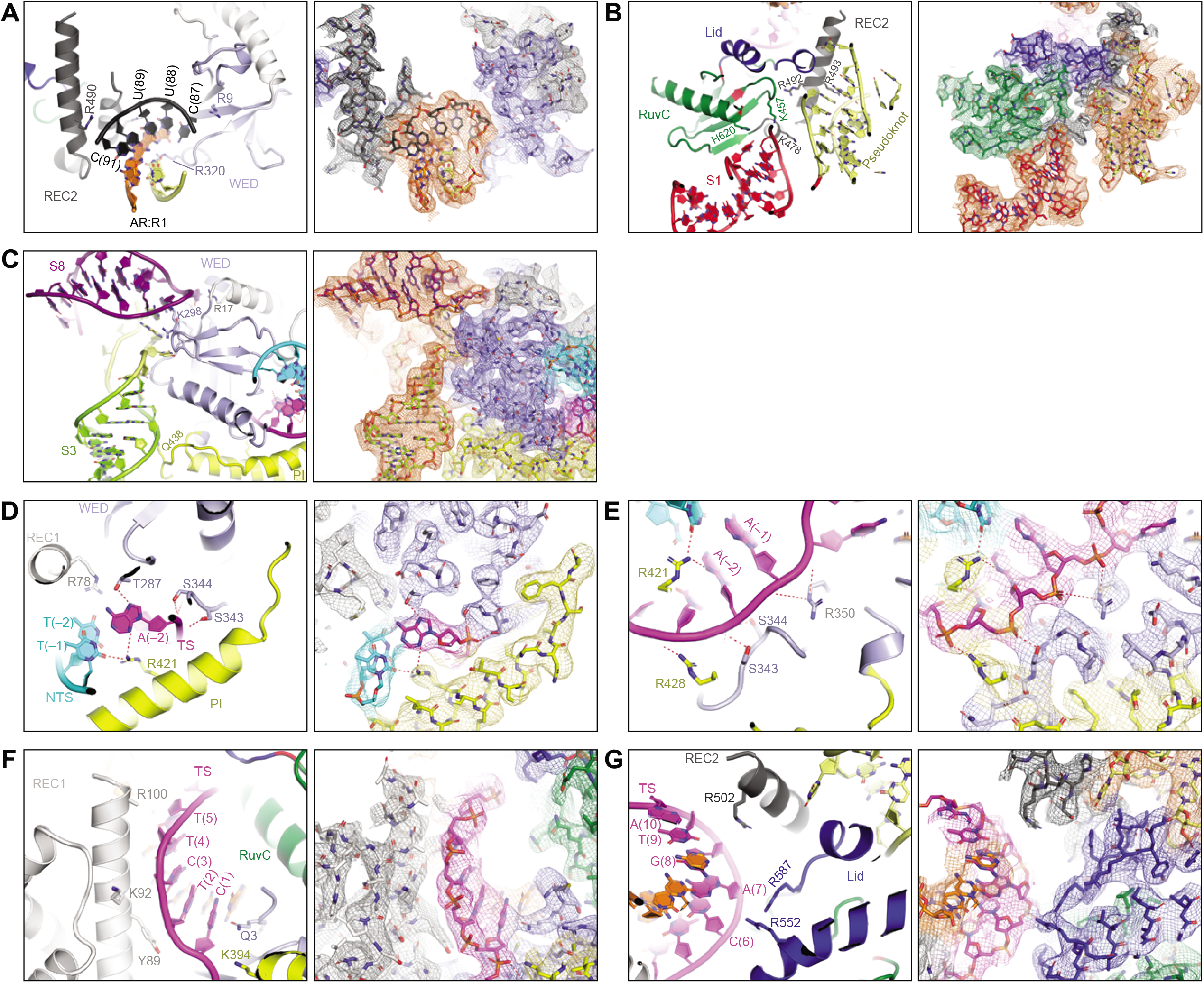
Interactions between Cas12k and bound nucleic acids. **(A–C)** Contacts between Cas12k and sgRNA. The AR:R1 duplex is clamped between the N-lobe and C-lobe of Cas12k **(A)**. S1 and pseudoknot contact the C-lobe **(B)**, whereas S3 and S8 contact the N-lobe of Cas12k **(C)**. **(D,E)** Interactions between Cas12k and the PAM duplex of target DNA. Left panels are in cartoon representation, while right panels are in stick presentation with cryo-EM density shown in mesh. Key residues involved in interactions are labeled. Interactions are indicated by red dashed lines. **(F)** Contacts between the N-lobe of Cas12k and the TS of the crRNA-target DNA heteroduplex. **(G)** Contacts between the C-lobe of Cas12k and the TS of the crRNA-target DNA heteroduplex.

**Figure S5.**
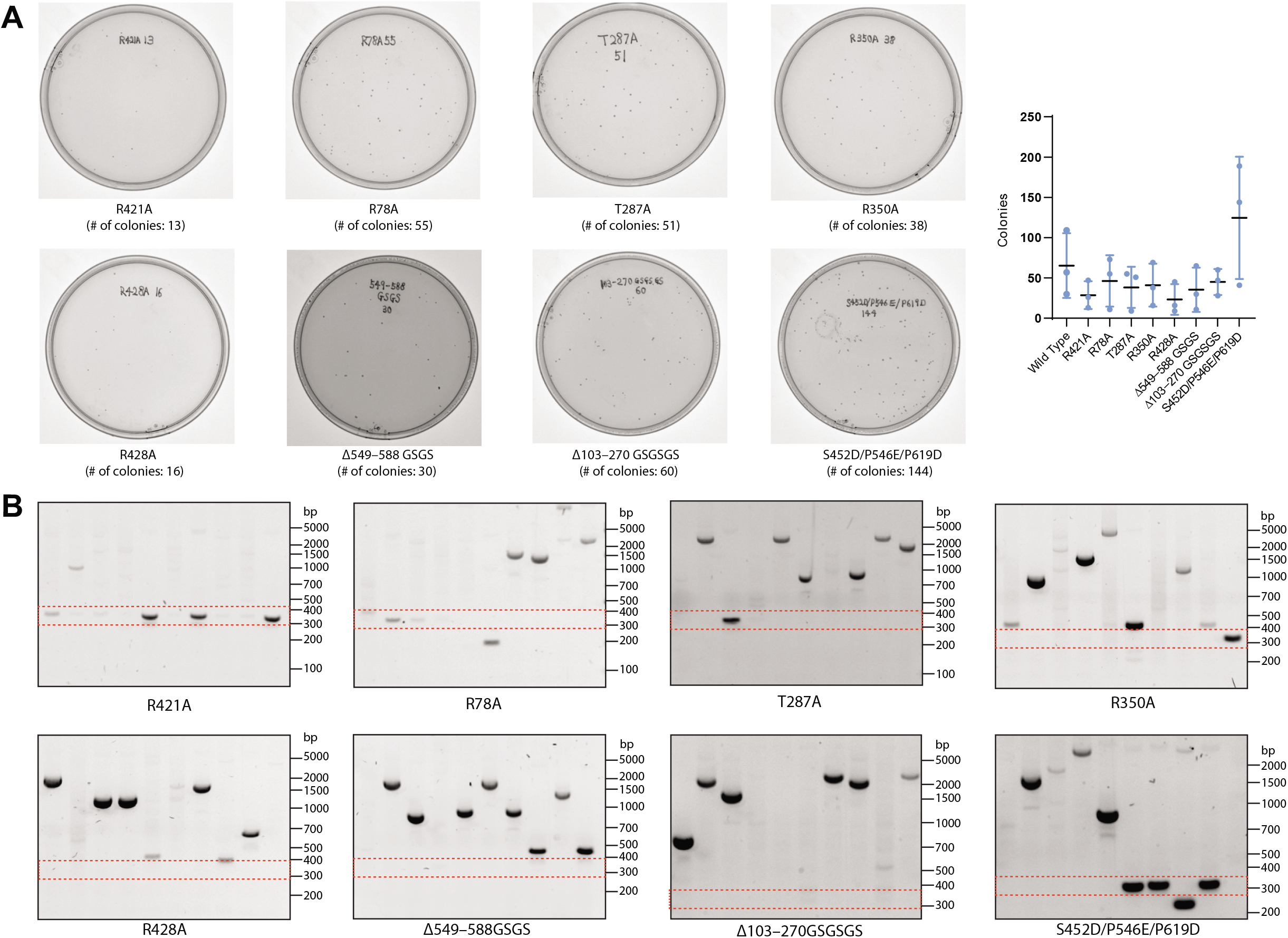
*In vitro* DNA transposition assay for Cas12k mutants. **(A)** LB agar plate showing colonies after transformation of each transposition reaction. Graph on the right shows mean±SD (n=3). **(B)** PCR results using purified plasmid as template. Ten colonies are randomly selected for plasmid extraction from each plate in **A**. Positions of expected PCR readout at ~350 bp are indicated by red dashed boxes. The plates and PCR results were from the same batch as those from **Fig. S1E,F**.

**Figure S6.**
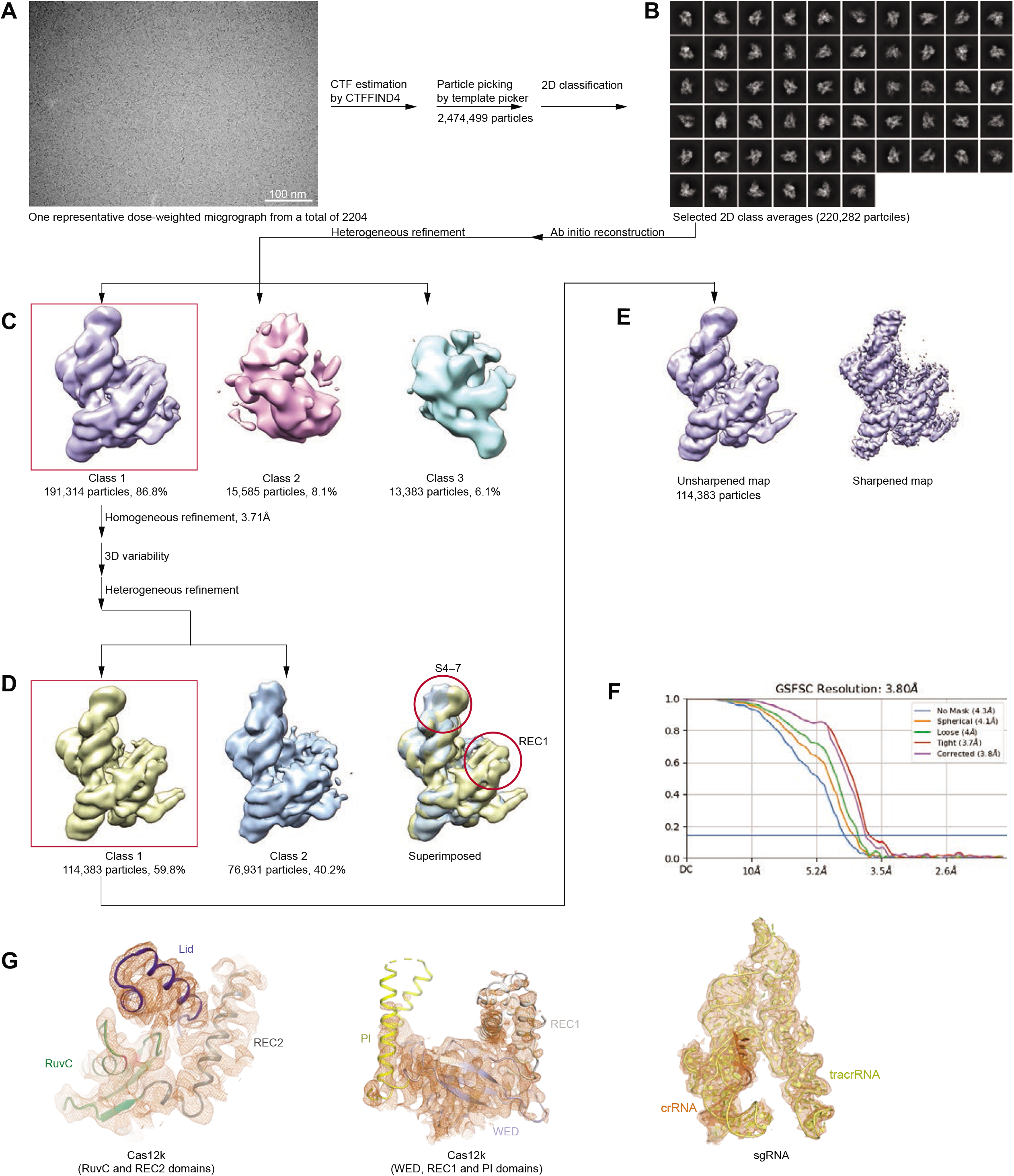
Cryo-EM data processing of the Cas12k–sgRNA binary complex. **(A)** A representative raw cryo-EM micrograph of the Cas12k–sgRNA complex from a total of 1849 micrographs. **(B)** Representative, good 2D class averages from a total of 100 images. **(C)** Three 3D reconstructions from heterogeneous refinement. **(D)** Supervised heterogenous refinement using two maps in 3D variability analysis as templates. Variable regions are indicated by red circles. **(E)** Homogeneous refinement of final particle set. Shown on the left is the unsharpened map, and on the right is the sharpened map. **(F)** Plots of the half-map FSC. **(G)** Fitting between the cryo-EM map of the Cas12k-sgRNA complex and the atomic model.

**Table S1. Sequence of RNAs and DNA oligonucleotides utilized in this study.**

## MATERIALS AND METHODS

### Protein expression and purification

Gene fragments of TnsB, TnsC, and TniQ were ordered from Integrated DNA Technologies (IDT) and cloned into bacterial expression plasmid pET28-MKH8SUMO (Addgene: #79526). The gene fragment for Cas12k was cloned into the bacterial expression plasmid pET-His6-StrepII-TEV LIC (Addgene: #29718). Cas12k, TnsB, and TniQ were expressed in *Escherichia coli* BL21(DE3) while TnsC was expressed in Rosetta™(DE3)pLysS (Novagen: #70956) containing a pLysS-tRNA plasmid. Cells were grown to OD_600_ =0.6 in Terrific Broth (TB) and protein expression was induced by adding 0.3 mM of IPTG followed by overnight incubation at 16°C. The cells were collected and resuspended in lysis buffer (50 mM Tris-HCl, pH□7.6, 500 mM NaCl, 5% glycerol) supplemented with 1 mM PMSF and 5 mM β-mercaptoethanol, and then disrupted by sonication. Cell lysate was clarified by centrifugation. The supernatant was loaded onto Ni-NTA resin. After extensive washing with lysis buffer supplemented with 30 mM imidazole, target proteins were eluted with lysis buffer supplemented with 250 mM imidazole. The His-SUMO tag of TnsB, TnsC, and TniQ and His-StrepII tag of Cas12k were removed by overnight digestion with TEV protease at 4°C. The protein was diluted with buffer containing 50 mM Tris-HCl pH 7.6, 200 mM NaCl, and 5% glycerol and loaded onto a Heparin column (GE Healthcare), eluted with a linear NaCl gradient (0.1 to 1M). After concentration, the proteins were further purified by size exclusion chromatography (SEC) over a Superdex 200 increase 10/300 GL column (Cytiva) in buffer containing 25 mM Tris-HCl (pH□7.6), 500 mM NaCl, 10% glycerol, and 1 mM DTT (0.5 mM EDTA was added to the buffer for TnsC). Fractions were concentrated and stored at −80°C.

To assemble the Cas12k–sgRNA binary complex, Cas12k proteins were incubated with sgRNA **(Table S1)** at a ratio of 1:1.15 at 37°C for 30 min in buffer A (25 mM Tris-HCl, pH□7.6, 150 mM NaCl, 2 mM DTT and 1 mM MgCl_2_). To reconstitute the Cas12k–sgRNA–target DNA ternary complex, Cas12k protein was incubated with sgRNA at 37°C for 30 min followed by the addition of target DNA synthesized from IDT **(Table S1)** at a ratio of 1:1.1.5:1.3. After 30 min, the mixture was subjected to SEC over a Superdex 200 column (Cytiva) equilibrated with buffer A for further purification.

### sgRNA preparation

sgRNAs were produced by *in vitro* transcription using the HiScribe T7 High Yield RNA synthesis kit (NEB) with PCR amplified gBlocks (IDT) as templates. sgRNAs were purified over a Resource-Q column (Cytiva) and eluted with a linear NaCl gradient (50 mM–1000 mM) in 25 mM Tris-HCl, pH 8.0. The eluted sgRNAs were concentrated and stored at −80°C

### Mutagenesis

Single amino acid mutations were introduced by the QuikChange site-directed mutagenesis method. Mutations with multiple amino acids were introduced by ligating inverse PCR-amplified backbone with mutations bearing DNA oligonucleotides via the In-Fusion Cloning Kit (ClonTech). All mutants were confirmed by Sanger sequencing.

### *In vitro* transposition assay

Donor plasmid (pDonor) and target plasmid (pTarget) were gifts from Feng Zhang (Addgene #127924 and #127926, respectively). *In vitro* transposition reaction was conducted as previously described unless otherwise stated. All proteins were diluted to 2 μM with 25 mM Tris-HCl, pH 8.0, 500 mM NaCl, 1 mM EDTA, 1 mM DTT, and 25% glycerol. 50 nM of each proteins, 600 nM sgRNA, 20 ng pTarget, and 100 ng pDonor were added sequentially to the reaction buffer containing 26 mM HEPES pH 7.5, 4.2 mM Tris-HCl pH 8.0, 2.1 mM DTT, 0.05 mM EDTA, 0.2 mM MgCl_2_, 28 mM NaCl, 21 mM KCl, 1.35% glycerol, 50 μg/mL BSA, and 2 mM ATP (final pH 7.5) to a total volume of 20 μL. Reactions were incubated at 30°C for 40 min before being supplemented with 25 mM MgOAc2 and incubated at 37°C for another 2 hours. 1 μL of the final products was taken out for direct PCR readout. The remaining sample was digested with 1 μL of Proteinase K (Thermo Fisher Scientific) at 37°C for 15 min before transformation into Stellar competent cells. Colonies were grown on kanamycin and chloramphenicol plates. Single colonies were randomly picked for plasmid preparation. After extraction, plasmids were analyzed by PCR, restriction enzyme (BamHI) digestion and sanger sequencing.

### Polymerase Chain Reaction (PCR)

Forward primer pTarget_F, reverse primer pDonor_R (**Table S1**), and *in vitro* transposition reaction product were mixed to a final volume of 25 μL for PCR reactions. Cycling conditions were as follows: 1 cycle, 94°C, 3 min; 35 cycles, 98°C, 10 s, 66.9°C, 15 s, 72°C, 8 s; 1 cycle, 72°C, 10 min. Plasmids extracted from single colonies were analyzed by PCR under cycling conditions as follows: 1 cycle, 98°C, 3 min; 35 cycles, 98°C, 10 s, 69.9°C, 15 s, 72°C, 12 s; 1 cycle, 72°C, 10 min.

### Electron Microscopy

Aliquots of 4 μL Cas12k–sgRNA binary complex (1 mg/mL) and Cas12k–sgRNA–dsDNA ternary complex (1 mg/mL) were applied to glow-discharged UltrAuFoil holey gold grids (R1.2/1.3, 300 mesh). The grids were blotted for 2 seconds and plunged into liquid ethane using a Vitrobot Mark IV. Cryo-EM data were collected with a Titan Krios microscope operated at 300 kV and images were collected using Leginon (Suloway et al., 2005) at a nominal magnification of 81,000x (resulting in a calibrated physical pixel size of 1.05 Å/pixel) with a defocus range of 0.8–2.0 μm. The images were recorded on a K3 electron direct detector in super-resolution mode at the end of a GIF-Quantum energy filter operated with a slit width of 20 eV. A dose rate of 20 electrons per pixel per second and an exposure time of 3.12 seconds were used, generating 40 movie frames with a total dose of ~ 54 electrons per Å^2^. Statistics for cryo-EM data are listed in **Table 1**.

### Image Processing

Movie frames were aligned using MotionCor2 (Zheng et al., 2017) with a binning factor of 2. The motion-corrected micrographs were imported into cryoSPARC (Punjani et al., 2017). Contrast transfer function (CTF) parameters were estimated using CTFFIND4 (Rohou and Grigorieff, 2015). A few thousand particles were auto-picked without template to generate 2D averages for subsequent template-based auto-picking. The auto-picked and extracted particles were processed for 2D classifications, which were used to exclude false and bad particles that fell into 2D averages with poor features. An initial reconstruction was done in cryoSPARC using 100,000 particles (Punjani et al., 2017). Heterogenous refinement was further performed to sort out different conformational heterogeneity. To further screen homogenous particles, 3D variance analysis (Punjani and Fleet, 2021) was performed and the resulting maps with different conformations (frame_000.mrc and frame_019.mrc) are used for supervised heterogenous refinement. The homogeneous dataset was used for final 3D refinement with C1 symmetry, resulting in 3.65 Å resolution from 183,870 particles.

The Cas12k–sgRNA binary complex dataset were processed in a similar way as the ternary complex. 114,383 particles were selected for a final reconstruction at 3.80 Å resolution. Cryo-EM image processing is summarized in **Table 1**.

### Model building, refinement, and validation

*De novo* model building of the Cas12k–sgRNA–target DNA structure was performed manually in COOT (Emsley et al., 2010) guided by secondary structure predictions from PSIPRED (Jones, 1999) of Cas12k protein and structure prediction of sgRNA by RNAComposer (Biesiada et al., 2016). Refinement of the structure models against corresponding maps were performed using the *phenix.real_space_refine* tool in Phenix (version 1.19.2) (Afonine et al., 2018). For the Cas12k–sgRNA complex, the structure model of the Cas12k–sgRNA–target-DNA complex was fitted into the cryo-EM map with models for target DNA deleted. The model is adjusted by all-atom refinement in COOT with self-restrains. The resultant model was refined against the corresponding cryo-EM map using the *phenix.real_space_refine* tool in Phenix.

### Structure-based sequence alignment

PROMALS3D program (Pei et al., 2008) was used to align the sequences of Cas12k and Cas12f based on structure. The alignment diagram was plotted using ESPript (Robert and Gouet, 2014). Sequence identities and similarities were calculated using Sequence Manipulation Suite (Stothard, 2000). Root-mean-square deviation (RMSD) of the Cα atomic was calculated using the *cealign* command in PyMOL.

### Structural visualization

Figures were generated using PyMOL and UCSF Chimera (Pettersen et al., 2004).

## References

Afonine, P.V., Poon, B.K., Read, R.J., Sobolev, O.V., Terwilliger, T.C., Urzhumtsev, A., and Adams, P.D. (2018). Real-space refinement in PHENIX for cryo-EM and crystallography. Acta Crystallogr D Struct Biol 74, 531–544.

Biesiada, M., Purzycka, K.J., Szachniuk, M., Blazewicz, J., and Adamiak, R.W. (2016). Automated RNA 3D Structure Prediction with RNAComposer. Methods Mol Biol 1490, 199–215.

Dong, D., Ren, K., Qiu, X., Zheng, J., Guo, M., Guan, X., Liu, H., Li, N., Zhang, B., and Yang, D. (2016). The crystal structure of Cpf1 in complex with CRISPR RNA. Nature 532, 522.

Emsley, P., Lohkamp, B., Scott, W.G., and Cowtan, K. (2010). Features and development of Coot. Acta Crystallogr D Biol Crystallogr 66, 486–501.

Faure, G., Shmakov, S.A., Yan, W.X., Cheng, D.R., Scott, D.A., Peters, J.E., Makarova, K.S., and Koonin, E.V. (2019). CRISPR-Cas in mobile genetic elements: counter-defence and beyond. Nat Rev Microbiol 17, 513–525.

Gao, P., Yang, H., Rajashankar, K.R., Huang, Z., and Patel, D.J. (2016). Type V CRISPR-Cas Cpf1 endonuclease employs a unique mechanism for crRNA-mediated target DNA recognition. Cell research 26, 901.

Gratz, S.J., Ukken, F.P., Rubinstein, C.D., Thiede, G., Donohue, L.K., Cummings, A.M., and O’Connor-Giles, K.M. (2014). Highly specific and efficient CRISPR/Cas9-catalyzed homology-directed repair in Drosophila. Genetics 196, 961–971.

Halpin-Healy, T.S., Klompe, S.E., Sternberg, S.H., and Fernandez, I.S. (2020). Structural basis of DNA targeting by a transposon-encoded CRISPR-Cas system. Nature 577, 271–274.

Huang, X., Sun, W., Cheng, Z., Chen, M., Li, X., Wang, J., Sheng, G., Gong, W., and Wang, Y. (2020). Structural basis for two metal-ion catalysis of DNA cleavage by Cas12i2. Nat Commun 11, 5241.

Jia, N., Xie, W., de la Cruz, M.J., Eng, E.T., and Patel, D.J. (2020). Structure-function insights into the initial step of DNA integration by a CRISPR-Cas-Transposon complex. Cell Res 30, 182–184.

Jones, D.T. (1999). Protein secondary structure prediction based on position-specific scoring matrices. J Mol Biol 292, 195–202.

Klompe, S.E., Vo, P.L.H., Halpin-Healy, T.S., and Sternberg, S.H. (2019). Transposon-encoded CRISPR-Cas systems direct RNA-guided DNA integration. Nature 571, 219–225.

Li, Z., Zhang, H., Xiao, R., and Chang, L. (2020). Cryo-EM structure of a type I-F CRISPR RNA guided surveillance complex bound to transposition protein TniQ. Cell Res 30, 179–181.

Li, Z., Zhang, H., Xiao, R., Han, R., and Chang, L. (2021). Cryo-EM structure of the RNA-guided ribonuclease Cas12g. Nat Chem Biol.

Liu, J.J., Orlova, N., Oakes, B.L., Ma, E., Spinner, H.B., Baney, K.L.M., Chuck, J., Tan, D., Knott, G.J., Harrington, L.B., et al. (2019). CasX enzymes comprise a distinct family of RNA-guided genome editors. Nature 566, 218–223.

Liu, L., Chen, P., Wang, M., Li, X., Wang, J., Yin, M., and Wang, Y. (2017). C2c1-sgRNA Complex Structure Reveals RNA-Guided DNA Cleavage Mechanism. Mol Cell 65, 310–322.

Makarova, K.S., Wolf, Y.I., Alkhnbashi, O.S., Costa, F., Shah, S.A., Saunders, S.J., Barrangou, R., Brouns, S.J., Charpentier, E., Haft, D.H., et al. (2015). An updated evolutionary classification of CRISPR-Cas systems. Nat Rev Microbiol 13, 722–736.

Makarova, K.S., Wolf, Y.I., Iranzo, J., Shmakov, S.A., Alkhnbashi, O.S., Brouns, S.J.J., Charpentier, E., Cheng, D., Haft, D.H., Horvath, P., et al. (2020). Evolutionary classification of CRISPR-Cas systems: a burst of class 2 and derived variants. Nat Rev Microbiol 18, 67–83.

Mohanraju, P., Makarova, K.S., Zetsche, B., Zhang, F., Koonin, E.V., and van der Oost, J. (2016). Diverse evolutionary roots and mechanistic variations of the CRISPR-Cas systems. Science 353, aad5147.

Montano, S.P., Pigli, Y.Z., and Rice, P.A. (2012). The mu transpososome structure sheds light on DDE recombinase evolution. Nature 491, 413–417.

Moreno-Mateos, M.A., Fernandez, J.P., Rouet, R., Vejnar, C.E., Lane, M.A., Mis, E., Khokha, M.K., Doudna, J.A., and Giraldez, A.J. (2017). CRISPR-Cpf1 mediates efficient homology-directed repair and temperature-controlled genome editing. Nat Commun 8, 2024.

Nishimasu, H., Yamano, T., Gao, L., Zhang, F., Ishitani, R., and Nureki, O. (2017). Structural basis for the altered PAM recognition by engineered CRISPR-Cpf1. Molecular cell 67, 139–147. e132.

Park, J.-U., Tsai, A., Mehrotra, E., Petassi, M.T., Shan-Chi, H., Ke, A., Peters, J.E., and Kellogg, E.H. (2021). Structural basis for target-site selection in RNA-guided DNA transposition systems. bioRxiv.

Pei, J., Kim, B.H., and Grishin, N.V. (2008). PROMALS3D: a tool for multiple protein sequence and structure alignments. Nucleic Acids Res 36, 2295–2300.

Pettersen, E.F., Goddard, T.D., Huang, C.C., Couch, G.S., Greenblatt, D.M., Meng, E.C., and Ferrin, T.E. (2004). UCSF Chimera--a visualization system for exploratory research and analysis. J Comput Chem 25, 1605–1612.

Punjani, A., and Fleet, D.J. (2021). 3D variability analysis: Resolving continuous flexibility and discrete heterogeneity from single particle cryo-EM. J Struct Biol 213, 107702.

Punjani, A., Rubinstein, J.L., Fleet, D.J., and Brubaker, M.A. (2017). cryoSPARC: algorithms for rapid unsupervised cryo-EM structure determination. Nat Methods 14, 290–296.

Rice, P.A., Craig, N.L., and Dyda, F. (2020). Comment on “RNA-guided DNA insertion with CRISPR-associated transposases”. Science 368.

Robert, X., and Gouet, P. (2014). Deciphering key features in protein structures with the new ENDscript server. Nucleic Acids Res 42, W320–324.

Rohou, A., and Grigorieff, N. (2015). CTFFIND4: Fast and accurate defocus estimation from electron micrographs. J Struct Biol 192, 216–221.

Saito, M., Ladha, A., Strecker, J., Faure, G., Neumann, E., Altae-Tran, H., Macrae, R.K., and Zhang, F. (2021). Dual modes of CRISPR-associated transposon homing. Cell.

Shen, Y., Gomez-Blanco, J., Petassi, M.T., Peters, J.E., Ortega, J., and Guarné, A. (2021). Structural basis for DNA targeting by the Tn7 transposon. bioRxiv.

Shmakov, S., Abudayyeh, O.O., Makarova, K.S., Wolf, Y.I., Gootenberg, J.S., Semenova, E., Minakhin, L., Joung, J., Konermann, S., Severinov, K., et al. (2015). Discovery and Functional Characterization of Diverse Class 2 CRISPR-Cas Systems. Mol Cell 60, 385–397.

Skelding, Z., Sarnovsky, R., and Craig, N.L. (2002). Formation of a nucleoprotein complex containing Tn7 and its target DNA regulates transposition initiation. EMBO J 21, 3494–3504.

Sorek, R., Lawrence, C.M., and Wiedenheft, B. (2013). CRISPR-mediated adaptive immune systems in bacteria and archaea. Annu Rev Biochem 82, 237–266.

Stella, S., Alcón, P., and Montoya, G. (2017). Structure of the Cpf1 endonuclease R-loop complex after target DNA cleavage. Nature 546, 559.

Stella, S., Mesa, P., Thomsen, J., Paul, B., Alcon, P., Jensen, S.B., Saligram, B., Moses, M.E., Hatzakis, N.S., and Montoya, G. (2018a). Conformational Activation Promotes CRISPR-Cas12a Catalysis and Resetting of the Endonuclease Activity. Cell 175, 1856–1871 e1821.

Stella, S., Mesa, P., Thomsen, J., Paul, B., Alcón, P., Jensen, S.B., Saligram, B., Moses, M.E., Hatzakis, N.S., and Montoya, G. (2018b). Conformational Activation Promotes CRISPR-Cas12a Catalysis and Resetting of the Endonuclease Activity. Cell.

Stothard, P. (2000). The sequence manipulation suite: JavaScript programs for analyzing and formatting protein and DNA sequences. Biotechniques 28, 1102, 1104.

Strecker, J., Jones, S., Koopal, B., Schmid-Burgk, J., Zetsche, B., Gao, L., Makarova, K.S., Koonin, E.V., and Zhang, F. (2019a). Engineering of CRISPR-Cas12b for human genome editing. Nat Commun 10, 212.

Strecker, J., Ladha, A., Gardner, Z., Schmid-Burgk, J.L., Makarova, K.S., Koonin, E.V., and Zhang, F. (2019b). RNA-guided DNA insertion with CRISPR-associated transposases. Science 365, 48–53.

Strecker, J., Ladha, A., Makarova, K.S., Koonin, E.V., and Zhang, F. (2020). Response to Comment on “RNA-guided DNA insertion with CRISPR-associated transposases”. Science 368.

Suloway, C., Pulokas, J., Fellmann, D., Cheng, A., Guerra, F., Quispe, J., Stagg, S., Potter, C.S., and Carragher, B. (2005). Automated molecular microscopy: the new Leginon system. J Struct Biol 151, 41–60.

Swarts, D.C., and Jinek, M. (2019). Mechanistic Insights into the cis- and trans-Acting DNase Activities of Cas12a. Mol Cell 73, 589–600 e584.

Swarts, D.C., van der Oost, J., and Jinek, M. (2017). Structural Basis for Guide RNA Processing and Seed-Dependent DNA Targeting by CRISPR-Cas12a. Mol Cell 66, 221–233 e224.

Takeda, S.N., Nakagawa, R., Okazaki, S., Hirano, H., Kobayashi, K., Kusakizako, T., Nishizawa, T., Yamashita, K., Nishimasu, H., and Nureki, O. (2020). Structure of the miniature type V-F CRISPR-Cas effector enzyme. Mol Cell.

Teng, F., Cui, T., Feng, G., Guo, L., Xu, K., Gao, Q., Li, T., Li, J., Zhou, Q., and Li, W. (2018). Repurposing CRISPR-Cas12b for mammalian genome engineering. Cell Discov 4, 63.

Wang, B., Xu, W., and Yang, H. (2020). Structural basis of a Tn7-like transposase recruitment and DNA loading to CRISPR-Cas surveillance complex. Cell Res 30, 185–187.

Wu, D., Guan, X., Zhu, Y., Ren, K., and Huang, Z. (2017). Structural basis of stringent PAM recognition by CRISPR-C2c1 in complex with sgRNA. Cell Res 27, 705–708.

Xiao, R., Li, Z., Wang, S., Han, R., and Chang, L. (2021). Structural basis for substrate recognition and cleavage by the dimerization-dependent CRISPR-Cas12f nuclease. Nucleic Acids Res 49, 4120–4128.

Yamano, T., Nishimasu, H., Zetsche, B., Hirano, H., Slaymaker, I.M., Li, Y., Fedorova, I., Nakane, T., Makarova, K.S., and Koonin, E.V. (2016). Crystal structure of Cpf1 in complex with guide RNA and target DNA. Cell 165, 949–962.

Yamano, T., Zetsche, B., Ishitani, R., Zhang, F., Nishimasu, H., and Nureki, O. (2017). Structural basis for the canonical and non-canonical PAM recognition by CRISPR-Cpf1. Molecular cell 67, 633–645. e633.

Yan, W.X., Hunnewell, P., Alfonse, L.E., Carte, J.M., Keston-Smith, E., Sothiselvam, S., Garrity, A.J., Chong, S., Makarova, K.S., Koonin, E.V., et al. (2019). Functionally diverse type V CRISPR-Cas systems. Science 363, 88–91.

Yang, H., Gao, P., Rajashankar, K.R., and Patel, D.J. (2016). PAM-Dependent Target DNA Recognition and Cleavage by C2c1 CRISPR-Cas Endonuclease. Cell 167, 1814–1828 e1812.

Zetsche, B., Gootenberg, J.S., Abudayyeh, O.O., Slaymaker, I.M., Makarova, K.S., Essletzbichler, P., Volz, S.E., Joung, J., van der Oost, J., Regev, A., et al. (2015). Cpf1 is a single RNA-guided endonuclease of a class 2 CRISPR-Cas system. Cell 163, 759–771.

Zhang, H., Li, Z., Daczkowski, C.M., Gabel, C., Mesecar, A.D., and Chang, L. (2019). Structural Basis for the Inhibition of CRISPR-Cas12a by Anti-CRISPR Proteins. Cell Host Microbe 25, 815–826 e814.

Zhang, H., Li, Z., Xiao, R., and Chang, L. (2020). Mechanisms for target recognition and cleavage by the Cas12i RNA-guided endonuclease. Nat Struct Mol Biol.

Zheng, S.Q., Palovcak, E., Armache, J.P., Verba, K.A., Cheng, Y., and Agard, D.A. (2017). MotionCor2: anisotropic correction of beam-induced motion for improved cryo-electron microscopy. Nat Methods 14, 331–332.

